# Accounting for *cis*-regulatory constraint prioritizes genes likely to affect species-specific traits

**DOI:** 10.1101/2022.03.29.486301

**Authors:** Alexander L. Starr, David Gokhman, Hunter B. Fraser

## Abstract

Measuring allele-specific expression in interspecies hybrids is a powerful way to detect cis-regulatory changes underlying adaptation. However, it remains difficult to identify genes most likely to explain species-specific traits. Here, we outline a simple strategy that leverages population-scale allele-specific RNA-seq data to identify genes that have constrained *cis*- regulation within species yet show divergence between species. Applying this strategy to data from human-chimpanzee hybrid cortical spheroids, we identify signatures of lineage-specific selection on genes related to cellular proliferation, speech, and glucose metabolism. We also highlight cis-regulatory divergence in *CUX1* and *EDNRB* that may shape the unique trajectory of human brain development.

## Background

Changes in gene expression are thought to play a major role in the evolution of complex traits [1–4]. As a result, comparing gene expression between species can enable the identification of molecular changes underlying phenotypic divergence. However, obtaining accurate comparisons of gene expression between species is challenging due to confounding factors like age, environmental effects, differential cell type abundances, differences in developmental timing, and batch effects [5–7]. The use of interspecies hybrids overcomes these issues through the measurement of allele-specific expression (ASE) [8, 9]. In hybrids, the genomes of both species share the same nucleus and are exposed to identical environments, so there are no confounding environmental, batch, compositional, or developmental effects. This approach has been successfully applied in many species and advanced our understanding of the evolution of gene regulation and its role in establishing phenotypic variation [10–13]. Furthermore, the recent development of human-chimpanzee allotetraploid “hybrid” cells and organoids enables detailed, accurate quantification of differences in gene expression between humans and our closest living relatives [8,9,14].

Hybrids also enable the separation of *cis* and *trans* components of interspecies differences in gene expression [8, 9]. The *cis*-component is caused by differences in regulatory elements such as promoters or enhancers that only affect the expression of a nearby gene or genes on the same DNA molecule. The *trans*-component stems from changes in diffusible molecules such as transcription factors that can regulate gene expression throughout the genome. In hybrids the genomes of the two species are exposed to the same *trans* factors. As a result, allele-specific differences in gene expression can only be explained by *cis*-regulatory differences. In addition, using ASE to identify differentially expressed genes (referred to as AS-DEGs) enables elimination of many confounding factors (including the environmental, batch, compositional, and developmental timing effects mentioned above). This not only increases the signal-to-noise ratio, but also disentangles important, potentially evolutionarily significant gene-specific *cis*- regulation from broad *trans*-acting changes [4,15,16].

While the resulting list of AS-DEGs from hybrids is likely more accurate and isolates the *cis*- regulatory component, there are often thousands of AS-DEGs which makes it difficult to prioritize candidate genes and pathways that may have played a major role in evolution. Differential expression p-values and fold changes are commonly used to rank genes in comparative RNA-seq studies. However, large and significant fold changes may often result from low evolutionary constraint on their expression levels, as opposed to being under positive directional selection. These large fold changes in unconstrained genes (e.g. pseudogenes) could result in no or very limited phenotypic changes, since a lack of constraint implies a lack of phenotypic consequence of changes in expression. Therefore, a large and significant fold change alone is not sufficient to determine the importance of the gene in the evolution of the parental species. For example, consider a gene whose expression varies by two-fold between species. If this gene also varies by two-fold within each species individual members of the same species, it is unlikely to account for any species-specific phenotypes. In contrast, a gene that is under strong stabilizing selection—with little variation in expression within species but with a two-fold change between species–is more likely to have contributed to phenotypic divergence between species. Most studies of expression divergence between species do not include any comparison to within-species variation; the few exceptions have been limited by small sample sizes and the confounding factors inherent to any between-species comparison [17–20].

Here, we describe a method that leverages population-scale ASE data to approximate constraint on *cis*-regulation of gene expression. This method ranks AS-DEGs identified from interspecies hybrids in a way that is likely more able to prioritize adaptive, functionally significant changes. We apply this method to ASE data from human-chimpanzee hybrid cortical spheroids that recapitulate the gene expression patterns of the developing cerebral cortex. Using this dataset, we identify lineage-specific selection on the expression of genes related to speech, glucose metabolism, cellular proliferation, and glycan degradation [8, 9]. In addition, we highlight divergence in the expression of *CUX1* and *EDNRB* that may have played an important role in human brain evolution.

## Results

In a typical ASE pipeline, AS-DEGs are identified by comparing the RNA-seq read counts from the allele from species 1 and the allele from species 2 and ranked using a p-value for differential expression (Fig. 1A). Various enrichment tests and knowledge from the literature can then be used to identify interesting trends and prioritize candidate genes. However, these previous methods do not consider within-species variation in gene expression levels (Fig. 1B). If some genes have highly variable expression even within a single species, then differences of a similar (or smaller) magnitude between species are unlikely to explain species-specific traits (e.g. PDPR in Fig. 1B). Conversely, differential expression of genes with highly constrained expression in at least one species are more likely to be responsible for differences in organismal phenotypes between species. For example, ZNF331 and RPS16 have similar fold change magnitudes in human-chimpanzee hybrid cortical spheroids (Fig. 1C). However, the fold change for ZNF331 lies well within the distribution of fold changes between alleles in the human population whereas the fold changes for RPS16 are nearly outside the human population distribution (Fig. 1C). This indicates that the expression level of RPS16 is much more constrained and that its differential *cis*-regulation between human and chimpanzee is more likely to have phenotypic consequences.

**Figure 1.**
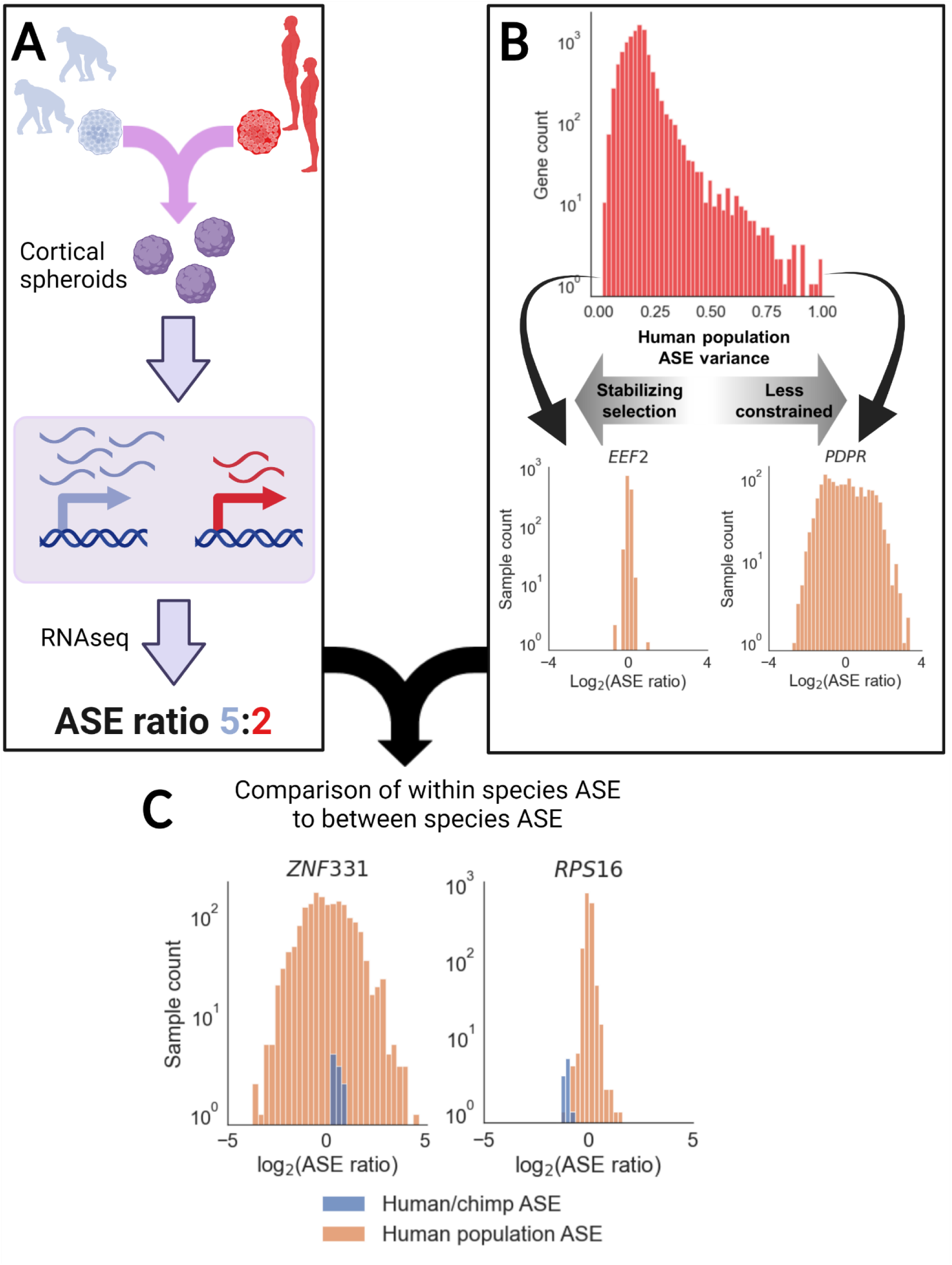
Outline of methodology: **A)** Outline of a typical ASE pipeline. Hybrids are generated and RNA-seq is used to determine the relative expression of each allele. The ASE ratio is computed as the ratio of species-specific read counts between the two alleles. **B)** The distribution of the variance in ASE ratio for each gene in the GTEx data. Insets in orange show two genes at the extreme ends (EEF2 with low variance suggesting strong stabilizing selection, and PDPR with high variance suggesting less constraint on gene expression). **C)** Schematic of incorporation of the interspecies ASE and population level ASE. ZNF331 has a wide range of ASE values, and the human-chimpanzee ASE is well within the population distribution whereas human-chimp ASE in RPS16 is on the edge of the population distribution indicating greater potential for functional relevance. For both **B** and **C**, only GTEx brain samples were used as this provided clearer illustrative examples, though results are similar using all GTEx samples (Fig 2A).

To systematically apply this concept, we compute the distribution of ASE in one population and use the Mann-Whitney U test to compare the population-level ASE distribution to the interspecies ASE distribution for that gene (see Methods) [21]. The p-value reflects how divergent the ASE of the gene is between species compared to its divergence within species. We then use those p-values to separately rank genes with DESeq2 false discovery rate (FDR) less than or equal to 0.1 and greater than 0.1 such that genes with FDR less than or equal to 0.1 are always ranked higher than genes with FDR greater than 0.1. This method prioritizes genes based on whether the interspecies difference in ASE is of greater magnitude than would be expected from the intra-species ASE distribution while minimizing false positives. Notably, ranking solely by the Mann-Whitney U Test p-value would potentially rank lowly expressed genes that are not significantly differentially expressed between species very highly, increasing the number of genes falsely identified as differentially expressed. This problem is eliminated by incorporating the DESeq2 FDR. The ranked list is then used in enrichment analyses and to identify of top candidates [22, 23].

We use a large publicly available data set, GTEx, to estimate ASE variance for every gene. The GTEx v8 data includes RNA-seq from 838 individuals and 54 tissues, a total of 15,253 samples in which ASE values have been previously estimated [21]. To minimize any allelic bias in the population distribution, we normalize the median to 1. The resulting rankings are robust to choosing either the reference or alternative allele as the numerator for the population distribution (spearman’s rho = 0.999 p < 10^-300^). Notably, we minimize any batch effects between GTEx and the human-chimpanzee hybrid dataset by focusing on ASE since direct comparisons of expression levels between data sets are not involved. For example, if some technical factor (e.g. sequencing platform or RNA isolation method) caused a gene to show a spurious 2-fold higher expression in GTEx samples compared to the hybrid data, this would cancel out in the GTEx ASE calculation.

First, we tested whether the variance in ASE in the human population is a reasonable proxy for constraint on gene expression. We compared the variance of the GTEx ASE distribution for each gene to its probability of haploinsufficiency score (pHI), a measure of sensitivity to a 50% reduction in gene dosage [24]. We observe a significant negative correlation (Spearman’s rho = -0.28, p < 10^-170^) between the two measures (Supp. Fig. 1) indicating that the variance of the population-scale ASE distribution provides a reasonable proxy for constraint on expression levels. This correlation remains significant when only five (as opposed to ten) reads from each allele for a gene in a sample are required for inclusion (Spearman’s rho = -0.27, p < 10^-163^). Furthermore, as pHI is computed from copy number variation, there is no circularity in reasoning when comparing pHI to the variance of the GTEx ASE distribution [24]. While this indicates that variation in ASE is sufficient to approximate evolutionary constraint on gene expression, the strength of this relationship should not be interpreted as a quantitative estimate of how well ASE represents constraint. Overall, the correlation with pHI indicates that within-species ASE variance contains useful information about evolutionary constraint on gene expression.

In addition, it is important to determine the extent to which sample heterogeneity may affect ASE variance and thereby impact our results. For example, GTEx contains data from a wide range of adult tissues and donors have variable sex and ancestry. To determine to what extent these factors affect our results, we divided the GTEx dataset along lines of sex (male/female), ancestry (African descent/European descent), and brain/non-brain tissues. In all cases, the results obtained were very highly correlated (spearman’s rho > 0.97, p < 10^-170^, Figure 2 A-C). This indicates that sample heterogeneity is unlikely to have major effects on our results and that aggregating across all the GTEx samples provides a reasonable measure of constraint on gene expression within the human population.

**Figure 2.**
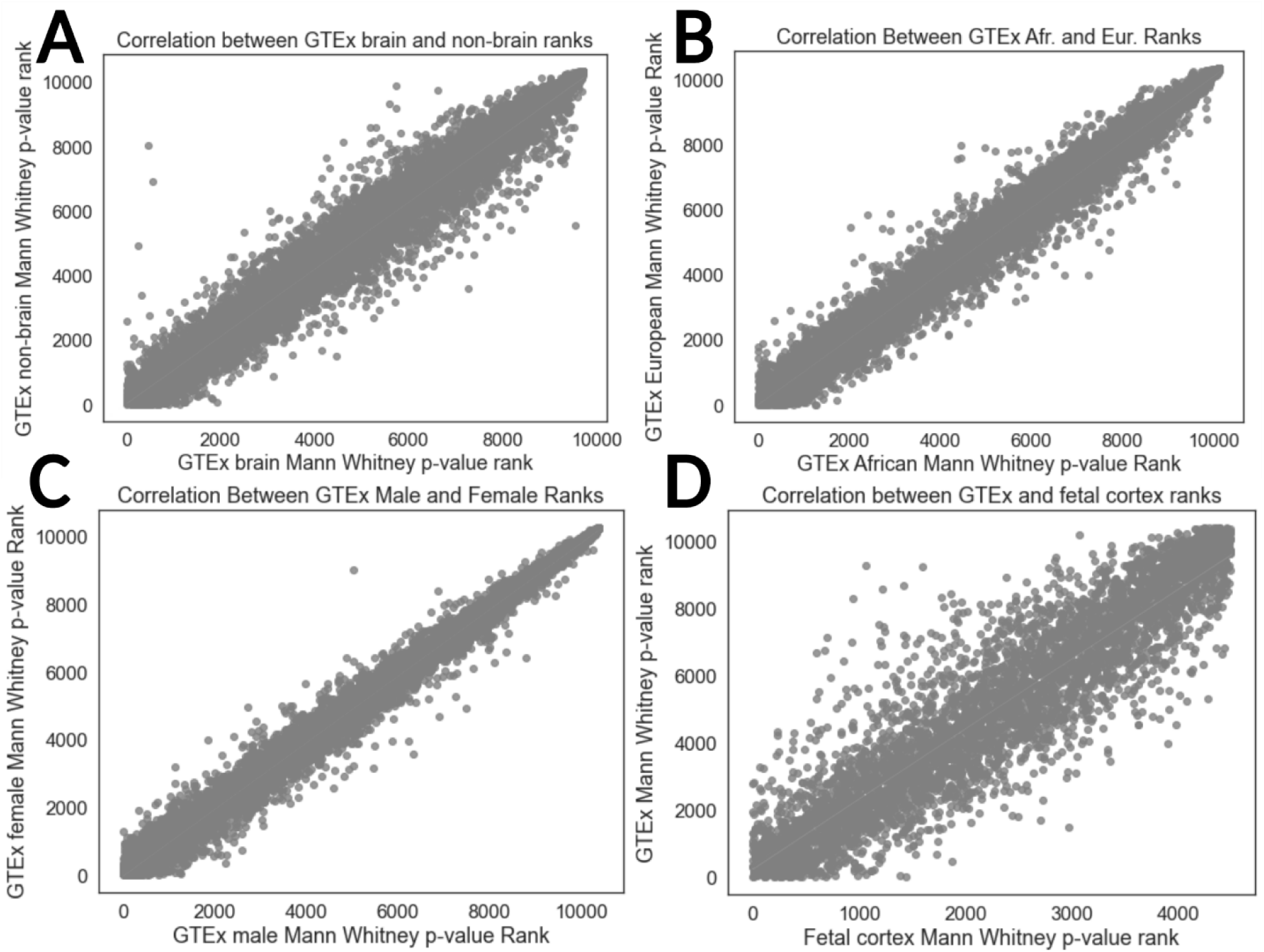
Influence of population differences on gene rankings: **A)** Comparison of ranks generated by our method derived from brain vs. non-brain samples. Spearman’s rho = 0.98 p < 10^-170^. **B)** Comparison of ranks using only GTEx subjects of African descent vs. subjects of European descent. Spearman’s rho = 0.98, p < 10^-170^. **C)** Comparison of ranks using only male vs. female GTEx subjects. Spearman’s rho = 0.99, p < 10^-170^. **D)** Comparison of ranks derived from GTEx (across all tissues) vs. data from fetal cortex and primary fetal neurons/neural progenitors. Spearman’s rho = 0.92, p < 10^-170^.

In addition, the wide range of adult tissues in GTEx may not accurately reflect gene expression constraint in cortical spheroids, which mimic fetal development [25, 26]. To test this, we used an ASE dataset generated from neural progenitors, neurons, and fetal cortical wall which resembles the developmental stage of cortical spheroids more closely than GTEx samples [27]. There is a strong correlation between the rankings generated by comparing to the fetal cortex dataset and GTEx (Spearman’s rho = 0.92, p < 10^-170^ for each of day 50, day 100, and day 150 after initial differentiation of cortical spheroids; Fig 2D). While it is likely that the ASE variance for some genes changes over development, these genes appear to be somewhat rare. Overall, these results (Fig 2) indicate that even though gene expression levels vary considerably across samples, ASE variance is robust to sample heterogeneity. This suggests that within- and between-species ASE values can be meaningfully compared even when they are not based on the same tissues or developmental timepoints. We therefore focused our analysis on the full GTEx data in order to maximize statistical power but verified results for specific genes in the fetal cortex data when appropriate.

To highlight differences between our method and the traditional method for ranking genes (Fig 1A vs. Fig 1C), we computed the difference in ranks between the two methods and used GSEAPY to identify gene sets enriched near the top or bottom of the resulting sorted list (referred to as the difference in ranks list, Fig 3A) [28]. A top-ranking gene would be one that is lowly ranked by the traditional method but highly ranked by the population comparison method. Many gene ontology (GO) categories related to cortical development were enriched near the top of the difference in ranks list including calcium and potassium channel activity, various transcription factor (TF) related terms, and cytokine activity (Fig 3B, C and Additional file 1). This is consistent with previous observations that TFs, ion channel subunits, and important players in canonical signaling pathways tend to have strongly constrained expression compared to other genes [24]. Many genes that drive the enrichment of the term “RNA polymerase II regulatory region sequence-specific DNA binding” are haploinsufficient and play key roles in neurodevelopment including *MEF2C*, *NEUROD2*, and *CUX1* (Supp. Fig. 2A) [29–31]. Differences in the expression of haploinsufficient genes are more likely to have phenotypic consequences than differences in the expression of genes for which loss of one copy has no clear phenotypic effect. In particular, there is strong evidence that both increases and decreases in CUX1 expression alter neurodevelopment in humans, which implies that the CUX1 expression divergence between humans and chimpanzees is likely to have phenotypic consequences (Box 1) [29, 32].

**Figure 3.**
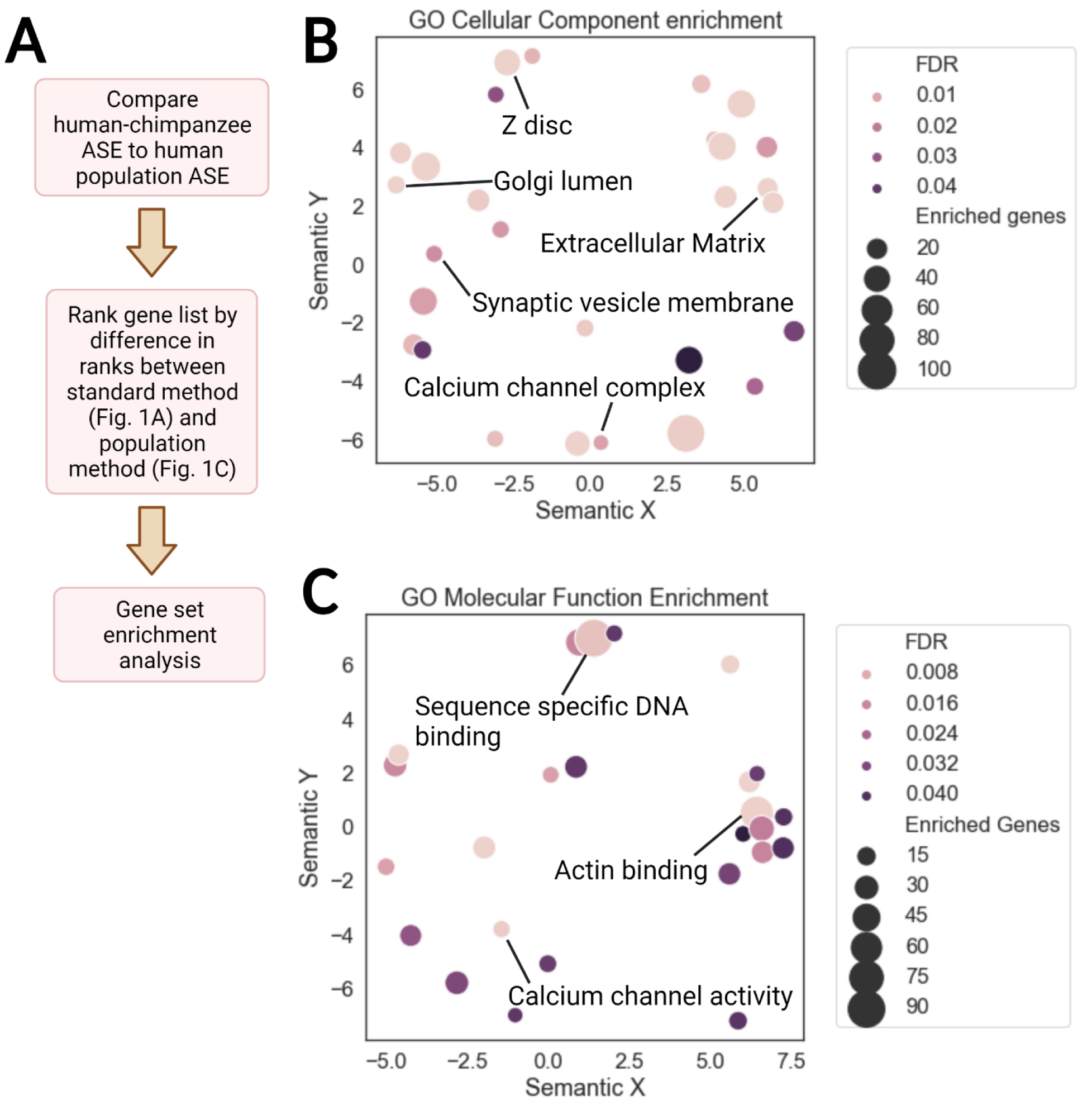
Enrichment summary for difference in ranks: **A)** Pipeline for gene set enrichment analysis. Gene set databases tested include Gene Ontology (GO) Cellular Component, GO Molecular Function, Kyoto Encyclopedia of Genes and Genomes (KEGG) human gene sets, Human Phenotype Ontology (HPO), and REACTOME [33–37]. Enrichment analysis is performed for each time point that cortical spheroids frozen for RNAseq (day 50, day 100, and day 150) separately and significant terms were aggregated with only one copy of redundant terms that were significant in multiple timepoints included. Only genes with sufficient reads in the human-chimpanzee cortical spheroid dataset and a sufficient number of samples in GTEx were included (see Methods), leaving approximately 10,000 genes in the list used for enrichment analysis. **B)** Summary of GO Cellular Component enrichments across all time points with false discovery rate (FDR) < 0.05. REVIGO in conjunction with a custom python script was used to generate the plot. The axes are derived from multidimensional scaling and measure semantic similarity, enabling removal of redundant GO terms and visualization of the similarity between GO categories. Each circle represents a GO term and circles near each other contain similar genes in the corresponding gene set. Labeled gene sets are generally those with the lowest FDR in a cluster of terms on the plot. The size of the circles indicates the number of genes that are driving the enrichment for that category. **C)** Summary of GO Molecular Function enrichments across all time points with FDR < 0.05.

### Box 1

Recent work has linked a mutation in a human accelerated region (HAR) that likely increases *CUX1* expression to autism spectrum disorder, suggesting that human-derived changes in *CUX1* expression alter human behavior [32, 38]. *CUX1* is expressed at a lower level in humans than chimpanzees across time points in both hybrid and parental cortical spheroids (e.g. log_2_ fold-change of -0.93, FDR < 0.005 in parentals, mean log_2_ fold-change of -0.74, FDR < 0.023 in hybrids at day 150 of differentiation) and the per-sample ASE ratios are well outside the human fetal cortex population ASE distribution (Supp. Fig. 2 B-E). This reduced expression in humans is surprising considering that a recent massively parallel reporter assay (MPRA) found that the HAR linked to *CUX1* should increase expression in humans [38]. As expression is lower in humans, haploinsufficiency of *CUX1* might provide a reasonable model of the phenotypic consequences of this change. *CUX1* haploinsufficiency in humans leads to delayed development of speech and motor skills [29]. One aspect of this condition is that individuals with one functional *CUX1* allele often close the developmental gap over time (i.e. cognitive impairments and delays disappear with age) [29]. This “catch up” phenotype is very rare and may even be specific to *CUX1* haploinsufficiency [29]. Interestingly, humans develop more slowly than other great apes (known as neoteny) but eventually “catch up,” reminiscent of the *CUX1* phenotype. While changes in *CUX1* expression may have played a role in causing human neoteny, investigation of the development of layer II-III cortical neurons and behavior of *CUX1* haploinsufficient mice will be required to explore this further.

### Applying the Sign Test to Constrained Differentially Expressed Genes

While our method is designed to prioritize genes that are relevant to phenotypic differences between species, it does not, on its own, imply selection on changes in gene expression. On the other hand, the sign test is designed to detect lineage-specific selection based on systematic up- or down-regulation of genes that deviates from neutral expectation [39, 40]. One important requirement of the test is that the expression divergence of every gene being tested should be driven by independent genetic differences. For example, a single mutation in a *trans*-acting factor could cause a whole pathway to be down-regulated, but this would only count as one genetic difference. For this reason, hybrids are ideal for applying the sign test to gene expression, since *cis*-regulation of genes is typically independent (except in the case of some neighboring genes that share *cis*-elements, which can be accounted for by using a minimum distance threshold between genes). For example, in the human-chimpanzee hybrid, if a particular pathway contains significantly more genes with higher expression from the human allele than the chimpanzee allele (significance determined by a binomial test), that would provide evidence of selection on that pathway in the human and/or chimpanzee lineage. Similar to all tests of selection, the sign test cannot discern what the cause of the selective pressure is. For example, many *cis*-regulatory changes could be compensating for a change in a single *trans*-acting factor. Alternatively, changes in gene expression might cause a phenotypic change that is being selected for. In either case, the sign test provides evidence of lineage-specific selection on cis-regulation.

While our method (Fig 1C) does not require comparison to the traditional method (Fig 1A), it is still of interest to determine if any of the categories that contain genes with strong differences in ranking between our method and the traditional method show signatures of lineage-specific selection. We therefore applied the sign test to identify gene sets in which highly ranked genes (i.e. those with a difference in ranks between the population comparison method and the traditional method above the cutoff found by GSEAPY) show a systematic bias in directionality. Three gene sets are significant by the binomial test at an FDR cutoff of 0.1: *Neuronal Cell Body*, *Cytokine Activity* and *Cytoskeleton* (Additional file 2, FDR = 0.085 for all three terms). The *Cytokine Activity* category is dominated by genes that promote stem cell proliferation which may have consequences for the proliferation of human and chimpanzee radial glia (although we caution against overinterpretation of this result given the low fold changes of most genes in this category; Box 2).

#### Box 2

The *Cytokine Activity* category is enriched near the top of the difference in ranks list with 15 out of 16 genes driving the enrichment displaying chimpanzee-biased expression (Fig. 4A, B). It is dominated by genes that generally promote neural stem cell proliferation, which is unexpected considering the directionality bias and the higher proliferative capacity of human neural stem cells. *EDN1* had the strongest chimpanzee bias (mean log2 fold-change = -0.92, FDR < 0.025 at Day 150, Fig 4B). As *EDN1* primarily signals through *EDNRB* in the brain [41], we also investigated the expression of *EDNRB*. Surprisingly, *EDNRB* is one of the most strongly human-biased genes across all timepoints in both hybrid and parental cortical spheroids (mean log2 fold-change = 4.46, FDR < 0.005 at Day 150 in hybrids, log2 fold-change = 2.85, FDR < 0.0005 in parental samples). Human-chimpanzee ASE generally exceeds ASE found in human populations for *EDNRB* (Fig 4C; Mann Whitney U Test comparing EDNRB distribution to human population distribution p = 0.00027 for *EDNRB*), although this is not the case for *EDN1* (Fig 4D). The human-biased *EDNRB* expression and chimpanzee-biased *EDN1* expression is generally consistent across timepoints in hybrid cortical spheroids (Fig 4E, F).

Changes in *EDNRB* expression appear to be human-derived with respect to gorillas and macaques (although orangutans may have independently acquired similar expression to humans in early-stage brain organoids) (Supp. Fig. 3 A, B). We were unable to confidently determine whether changes in *EDN1* expression were human- or chimpanzee-derived using currently available data (data not shown). Next, we analyzed single-cell RNA-seq data generated from 1 month, 2 month, and 4 month-old human and chimpanzee brain organoids to identify the cell types driving increased EDNRB expression. We found that a previously identified radial glial cell (RGC) cluster was characterized by high *EDNRB* expression with non-zero expression in over 50% of cells [42]. Furthermore, this cluster had higher expression than any chimpanzee cluster (Mann-Whitney U test, p < 10^-16^, Supp. Fig. 4-5). Overall, our results suggest that a subpopulation of human radial glia have much higher *EDNRB* expression than chimpanzee radial glia.

*EDNRB* haploinsufficiency reduces proliferation of cerebellar granule precursor cells and chemical inhibition of *EDNRB* signaling reduces proliferation of mouse radial glia [43, 44]. Based on this, the change in *EDNRB* expression may have promoted human brain expansion by increasing the proliferation of the subpopulation of radial glia that express *EDNRB*. In addition, a recent study identified a population of caudal late interneuron progenitor (CLIP) cells marked by expression of *EDNRB* and *PTGDS* along with caudal ganglionic eminence markers [45]. As both *EDNRB* and *PTGDS* have strongly human-biased ASE it would be interesting to investigate if this population of cells exists in chimpanzee brain organoids and if it may have expanded in humans. The phenotypic implications of the higher *ENDRB* and *PTGDS* but lower *EDN1* expression in humans will be an exciting area for further research.

Thus far, we have primarily focused on the differences between our new method and the traditional method for ranking genes. Having established their differences, we now turn to analysis of results from our new approach (Fig. 1 and Methods). To examine human- and chimpanzee-biased genes separately, we sorted the list so that highly ranked genes with human-biased expression are at the top of the list and highly ranked genes with chimpanzee-biased expression are at the bottom of the list. In effect, this results in a test for directionally-biased cis-regulatory divergence that exceeds the cis-regulatory variation among most human alleles present in the GTEx population. Enrichment testing with GSEAPY identified several enriched gene sets at an FDR cutoff of 0.25 (the cutoff suggested by the GSEA authors) including *Other glycan degradation* (human-biased), *Spastic Dysarthria* (human-biased), and *Gluconeogenesis* (chimpanzee-biased) (Fig 5 A-F, Supp. Table 3). In all three cases, there were zero genes showing expression bias in the other direction at an identical rank cutoff (p = 0.00012 for *gluconeogenesis*, p = 0.031 for *Other glycan degradation*, and p = 0.00195 for *Spastic Dysarthria* by binomial test), suggesting lineage-specific selection on genes with constrained expression in these gene sets [40]. Notably, the bias in gene expression found in the hybrids for these genes generally matched the bias in expression found in the parental cortical spheroids (Supp. Fig. 6A-C).

**Figure 5.**
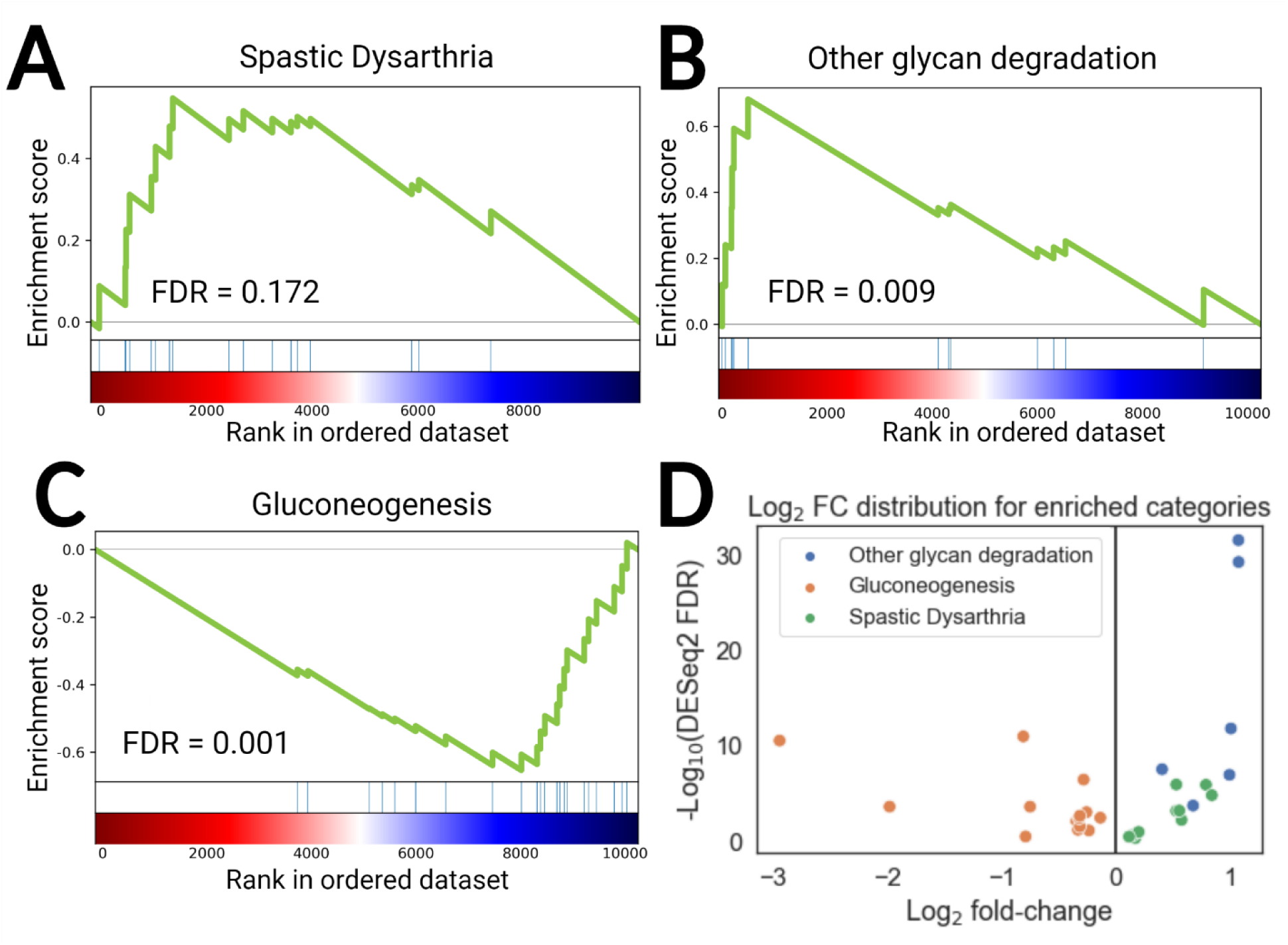
Evidence of lineage-specific selection: **A)** Summary of *Spastic dysarthria* enrichment (from Human Phenotype Ontology) [33, 37]. In A, B, and C, each blue line represents a gene in the gene set and the green curve is the cumulative enrichment score. **B)** Summary of *Other glycan degradation* enrichment (from KEGG). **C)** Summary of *Gluconeogenesis* enrichment (from REACTOME). **D)** Volcano plot summarizing of Log2 fold-changes for genes driving the enrichments for *Spastic Dysarthria*, *Other glycan degradation*, and *Gluconeogenesis*. All genes are human-biased for *Other glycan degradation* and *Spastic Dysarthria* whereas all genes are chimpanzee-biased for *Gluconeogenesis*.

All three of these enriched categories may influence human-specific phenotypes. Gluconeogenesis siphons oxaloacetate from the TCA cycle and eventually produces glucose [46]. Many of the gene expression changes driving the gluconeogenesis enrichment appear to be human-derived compared to other great apes (Supp. Fig. 7). Decreased gluconeogenesis in the human lineage would likely enable increased flux through other anabolic pathways and the TCA cycle possibly increasing the availability of oxaloacetate for anabolic pathways that promote proliferation. Spastic dysarthria is a condition in which patients speak in a characteristic slow, regular, monotone manner [47]. Loss of function of genes in this category are associated with spastic dysarthria and show systematic human bias, which may be connected to the human capacity for speech (Fig 5D). Finally, six genes that were ranked very highly by our method and all have DESeq2 FDR < 0.1 drive the “Other glycan degradation” enrichment. Interestingly, loss of function of three of the six human-biased glycan degradation genes (MANBA, MAN2B2, and MAN2B1) is associated with intellectual disability [48–50].

**Figure 4.**
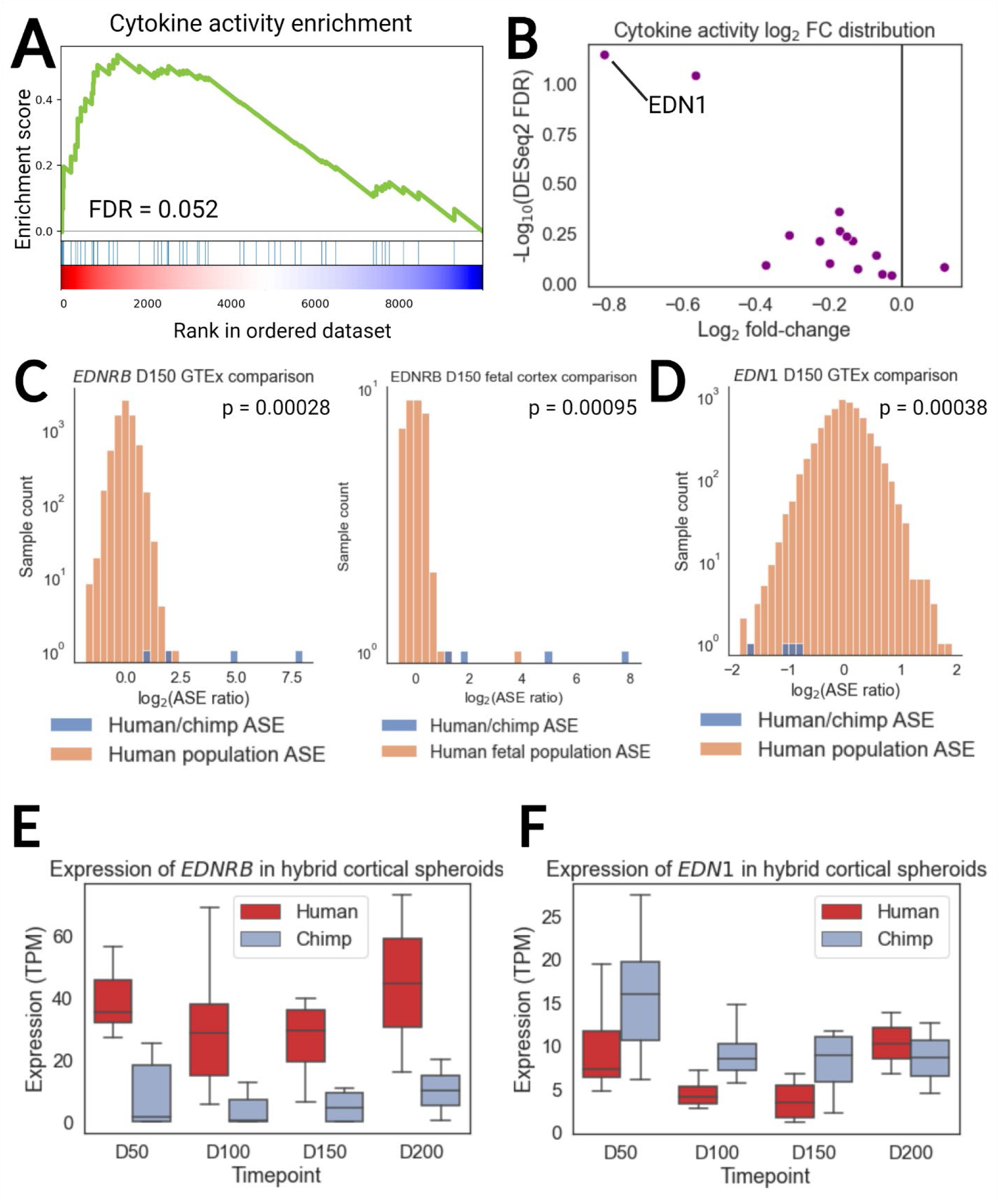
Changes in expression of cytokine activity genes: A) Summary of Cytokine Activity enrichment. Each blue line represents a gene in the gene set and the green curve is the cumulative enrichment score. Top ranking genes could be human or chimpanzee-biased in expression. B) Volcano plot showing expression levels of genes driving Cytokine Activity enrichment. The log_2_ fold-change (log_2_ FC, which refers to the value computed by DESeq2) is chimpanzee-biased for 15 out of 16 genes. Mean log_2_ FC between chimpanzee-referenced and human-referenced log_2_ FC values is shown. Negative log_2_ FC indicates chimpanzee-biased expression. EDNRB does not appear because it is a cytokine receptor rather than a cytokine. C) Comparison of human-chimpanzee EDNRB ASE to within-human ASE from GTEx and fetal cortical samples. Raw ASE ratios (as opposed to the value derived from DESeq2) are indicated by “ASE Ratio”. The human-chimpanzee ASE is significantly outside of the human ASE distribution. P-values are from the Mann-Whitney U Test comparing the distribution of human population ASE to human-chimpanzee ASE. D) Comparison of human-chimpanzee EDN1 ASE to within-human GTEx ASE. Fetal cortical ASE is not shown due to insufficient data. P-value is the same as in C. E) ASE of EDNRB across timepoints in cortical spheroids. Expression from the human allele is consistently higher than expression from the chimpanzee allele. F) ASE of EDN1 across timepoints in cortical spheroids. Expression from the chimpanzee allele is consistently higher than expression from the human allele except at D200.

## Discussion

Here we presented a method incorporating population-scale ASE data as a proxy for constraint on expression. This ranking method helps reveal candidate genes and signatures of selection that may explain phenotypic differences between humans and chimpanzees. The test is based on the logic of comparing within- to between-species variation, similar in spirit to the Hudson-Kreitman-Aguade test although it is based on variation in ASE rather than protein-coding sequences [51]. Although the phenotypic consequences of these differences remain to be determined, our finding of polygenic lineage-specific selection on several gene sets suggests that these changes must have some phenotypic effects in order to be under natural selection. Collectively, these findings more than double the number of known cases of lineage-specific polygenic selection on gene expression between humans and chimpanzees (the two previous examples being Hedgehog signaling and astrocyte-related genes) [8, 9].

Importantly, the strong correlation between the GTEx brain vs. non-brain rankings (Fig. 2A) and GTEx vs. fetal cortex rankings (Fig. 2D) suggests that comparison to the GTEx ASE distribution can be meaningfully compared with between-species ASE measured in diverse cell types and organoids and that comparison to the GTEx population distribution will be useful for other cell types and organoids. The method can also be applied to any species with sufficient gene expression data, e.g. comparing ASE in *Arabidopsis* interspecies hybrids to ASE within *A. thaliana* [52].

## Conclusion

We outlined a strategy that uses allele-specific expression data from interspecies hybrids and population-scale studies to prioritize genes that are more likely to impact species-specific traits and applied this method to data from human-chimpanzee cortical spheroids. Our findings provide opportunities for targeted follow-up experiments and increase our understanding of how polygenic selection has shaped human and chimpanzee evolution. Overall, we anticipate that our method will become a useful tool for identifying functionally significant gene expression changes between species, and will contribute to our understanding of how gene expression drives phenotypic diversification.

## Methods

### Read Alignment and RNA-seq Data Processing

Data from hybrid cortical spheroids was mapped as previously described [9]. Briefly, Hornet, a rewritten version of WASP, was used in conjunction with a curated list of human-chimpanzee SNPs and indels to correct for mapping bias. Reads for every sample were aligned to both the human and chimpanzee genomes and the log_2_ fold-change from both alignments was compared. Any genes with log_2_ fold-change that differed by greater than 1 were removed. We used the ASE log_2_ fold-change (log_2_ FC) values available in the supplemental tables of Agoglia et al. Although this dataset is restricted to hybrids from two humans and two chimpanzees, previous work has shown that interspecies differences dominate over differences between populations within a species so we expect that our results generalize well. It also contains multiple independently derived hybrid lines and independent differentiations, reducing confounding by technical differences. Throughout, for hybrids the mean log_2_ FC between human genome mapped and chimpanzee genome mapped reads is stated as well as the highest p-value. Additional data was downloaded from GSE127898, GSE106245, GSE153076, and phs000755.v2.p1 and mapped separately for each dataset to the respective species’ genome (PanTro6 for chimpanzee, hg38 for human, mmul10 for rhesus macaque, Gorgor6 for gorilla, and PonAbe3 for orangutan) [53–56]. We used STAR v2.5.4 with arguments: -outSAMattributes MD NH -outFilterMultimapNmax 1 -sjdbGTFfile -sjdbOverhang N where N is 1 less than the read length used for each respective dataset [57]. For paired end reads, we used Picard to remove duplicates with argument: DUPLICATE_SCORING_STRATEGY = RANDOM [58]. We used HT-Seq with the following arguments: -t exon -i gene_name -m intersection-strict -r pos to count reads overlapping gene bodies [59]. Transcripts per million (TPM) was computed as previously described [60]. We used the likelihood ratio test in DESeq2 to test for differential expression in the downloaded datasets with RIN and sex included as covariates for the Khrameeva et al. dataset [23, 61]. We binarized RIN values as high if greater than or equal to 7.5 and low otherwise as we do not generally expect the expression level of genes to scale linearly with RIN.

### Comparison of Population and Interspecies ASE Distributions

GTEx data was downloaded from https://www.gtexportal.org/home/datasets and the fetal cortex ASE data was kindly provided by the Stein laboratory [21, 27]. GTEx contains data from 838 individuals and the data from the Stein laboratory was generated from approximately 235 individuals. To preprocess the GTEx data, we split the file into two files each containing read counts from 1 of the alleles with a custom R script. As cortical spheroids are a mixture of different cell types including neural progenitors and immature neurons, we pooled fetal cortical wall, neural progenitor, and neuron counts per individual in the fetal cortex dataset. For each gene in each sample, we added one count to each gene (to prevent division by zero) and computed the ASE ratio as the ratio of counts from allele 1 (the reference) to counts from allele 2. The distribution for each gene was then normalized so that the median was 1. This normalization ensures that the Mann-Whitney U-test p-values only take into account the variance in allelic expression in the human population and are not confounded by consistently higher/lower expression from a particular allele. Notably, flipping the sign of each value in the GTEx ASE distribution had minimal effect on the rankings (Spearman’s rho = 0.999, p < 10^-300^ for day 50, day 100, and day 150 after the beginning of cortical spheroid differentiation) supporting the efficacy of the correcting the median to 1 in isolating the variance of the human population expression distribution. All samples with at least 10 counts (not including the single added count) from each allele for a sample were included in the ASE population distribution. Notably the rankings and our results are robust to requiring at least 5 counts from each allele instead of 10 (Spearman’s rho = 0.996, p < 10^-300^). To filter out genes that are lowly expressed in cortical spheroids, we removed genes with an average number of counts from the chimpanzee and human alleles less than 25 (i.e. mean of human and chimpanzee read count less than 25 and mean of chimpanzee read count less than 25). In addition, we filtered out any genes showing mapping bias (listed in the supplemental tables of Agoglia et al.) as well as genes on chromosomes 18 and 20 as parts of these chromosomes were duplicated in some cortical spheroid samples. Previous work has shown that these structural changes have minimal effect on the computation of ASE values for genes outside the duplicated region [9]. After filtering, we computed the interspecies ASE distribution in similar manner to the population ASE distribution (i.e. by taking the ratio of the counts from the human allele to the ratio of the counts of the chimpanzee allele). However, we did not require 10 counts from each allele and did not normalize the medians. We did not require 10 counts from each allele because we expect extreme differences in expression to be relatively common in between species comparisons. We compared the log2(ASE Ratio) interspecies distribution to the population distribution using the Mann-Whitney U Test (a nonparametric test robust to the distribution of data) and used the resulting p-values to rank genes as described below.

### Generation of Gene Rankings and Enrichment Analysis

First, we ranked genes by the Mann-Whitney U Test p-value with the lowest p-value receiving the highest rank. To reduce false positives at the top of the list, we separately ranked genes with DESeq2 FDR less than or equal to 0.1 and greater than 0.1. We then concatenated the list so that genes with FDR less than or equal to 0.1 were always ranked higher than genes with FDR greater than 0.1 (referred to as the MWU ranking). This consensus gene ranking was then used in GSEAPY preranked with the rankings used as the score that GSEAPY uses to sort the list. All results highlighted in the text replicated when using 0.05 as a cutoff instead of 0.1. We used REVIGO in conjunction with a custom python script to generate the plots shown in figure 2 [62].

To compare to the traditional method, we also ranked genes by the DESeq2 derived FDR and used that in GSEAPY preranked (referred to as the DESeq2 ranking). To highlight differences between the two methods, we computed the difference in ranks between the two methods by subtracting the DESeq2 ranking from the MWU ranking and sorting the list on those rankings for use in GSEAPY. In this context, highly ranked genes are likely those that show relatively mild gene expression changes but have more constrained expression.

We next performed the expression sign test. First, we generated a list of all gene sets across all tested ontologies that were nominally enriched at an FDR of 0.25 (using the FDR from GSEAPY preranked) in at least one time point and that had greater than 10 genes driving the enrichment. To avoid testing the same gene set multiple times, we only tested each gene set at the timepoint that had the lowest GSEAPY FDR. We used the average of the DESeq2 log_2_ FC from mapping to the human allele and from mapping to the chimpanzee allele as input for the binomial test to identify gene sets with significantly more human-biased or chimpanzee-biased changes than expected by chance. This log_2_ FC was generated by comparing the reads from each species’ allele in the cortical spheroid data. For example, if a gene set had 8 human-biased and 3 chimpanzee-biased genes, then the binomial test was used with k = 8, n = 11, and p = 0.5. We considered any gene set with Benjamini-Hochberg corrected FDR < 0.1 to be significant.

Finally, we ranked the genes using a signed version of the MWU ranking. More specifically, genes were effectively ranked by the log_10_(MWU p-value) multiplied by the sign of mean DESeq2 log_2_ fold-change so that top ranked genes with negative L2FC are at the bottom of the list and top ranked genes with positive L2FC are at the top of the list. This ranking was then used in GSEAPY preranked. All statistical tests (Mann Whitney U Test, Binomial Test, correlations) were performed in python using the implementation in scipy.

For enrichment testing, we tested gene sets from the Gene Ontology Cellular Component and Molecular Function categories, the Human Phenotype Ontology, KEGG, and REACTOME using the same version as in Gokhman et. al [8,33–37]. Regardless of which ranking was used, we used GSEAPY preranked with the following arguments: processes=4, permutation_num=1000, seed=6, min_size = 10, max_size = 300 to test for enrichment [28]. Following the authors suggestion, we considered any category with an FDR below 0.25 to be nominally enriched [28]. We required that ASE data be available from at least 50 individuals in GTEx for a gene to be included in the ranking. We conducted gene set enrichment analysis with the four different rankings described above.

### Single Cell RNA-seq Data Processing and Analysis

Single cell data from human and chimpanzee organoids and associated metadata were downloaded from E-MTAB-7552 [42]. We used SCANPY to read in the counts matrix and filter the data so that only data from 1 month, 2 month, and 4 month old organoids remained [63]. We used a two-sided Mann Whitney U Test to compare *EDNRB* log2(counts per million) between the “RGC early 2” cluster and all chimpanzee clusters with and without cells with 0 *EDNRB* counts included. Mean counts by cell type and tissue for fetal human expression were downloaded from GSE156793 [64]. We compared cerebrum counts to all other organs except eye and cerebellum due to their similarity to the cerebrum and used a binomial test to determine if more tissues exhibited higher expression than brain than expected by chance.

## Declarations

### Ethics approval and consent to participate

Access to restricted data was approved by dbGaP.

### Consent for publication

Not relevant to our study.

### Availability of data and materials

No new data nor materials were generated during this study. The datasets supporting the conclusions of this article are available in GSE, dbGaP, or the supplemental material of Agoglia et al. 2021 (https://www.nature.com/articles/s41586-021-03343-3#Sec35), GSE127898 (https://www.ncbi.nlm.nih.gov/geo/query/acc.cgi?acc=GSE127898), GSE106245 (https://www.ncbi.nlm.nih.gov/geo/query/acc.cgi?acc=GSE106245), GSE153076 (https://www.ncbi.nlm.nih.gov/geo/query/acc.cgi?acc=GSE153076), and phs000755.v2.p1 (https://www.ncbi.nlm.nih.gov/projects/gap/cgi-bin/study.cgi?study_id=phs000755.v2.p1), phs002493.v1.p1 (link unavailable, data was received directly from the Stein lab). The GTEx data was downloaded from https://www.gtexportal.org/home/datasets (although the specific file that was downloaded is no longer available). The software used to perform the analysis in this paper is available at: https://github.com/astarr97/Z_Score. There is no restriction on the use of this code.

### Author contributions

HBF, DG, and ALS conceptualized the study. ALS performed all analysis, validation, visualization, and writing of software. ALS wrote the original manuscript and HBF and DG reviewed and edited the manuscript. All authors approved the publication of the manuscript.

## Acknowledgements

We thank Jason Stein and Nil Aygun for providing processed fetal cortex, neuron, and neural progenitor ASE calls. We thank members of the Fraser lab for helpful discussion. Illustrations were created with BioRender.

**Description of Additional file 1**

File name is Additional file 1.csv. Enrichment testing for difference between traditional and population-based method. This file contains the output from GSEAPY for enrichment testing as well as the dataset that was used for the enrichment analysis.

**Description of Additional file 2**

File name is Additional file 2.csv. Sign test on enriched terms for difference between traditional and population-based method. This file contains the results for the sign test applied to a subset (see Methods) of the significant terms from Additional file 1.

---

**Supplementary Figure 1.**
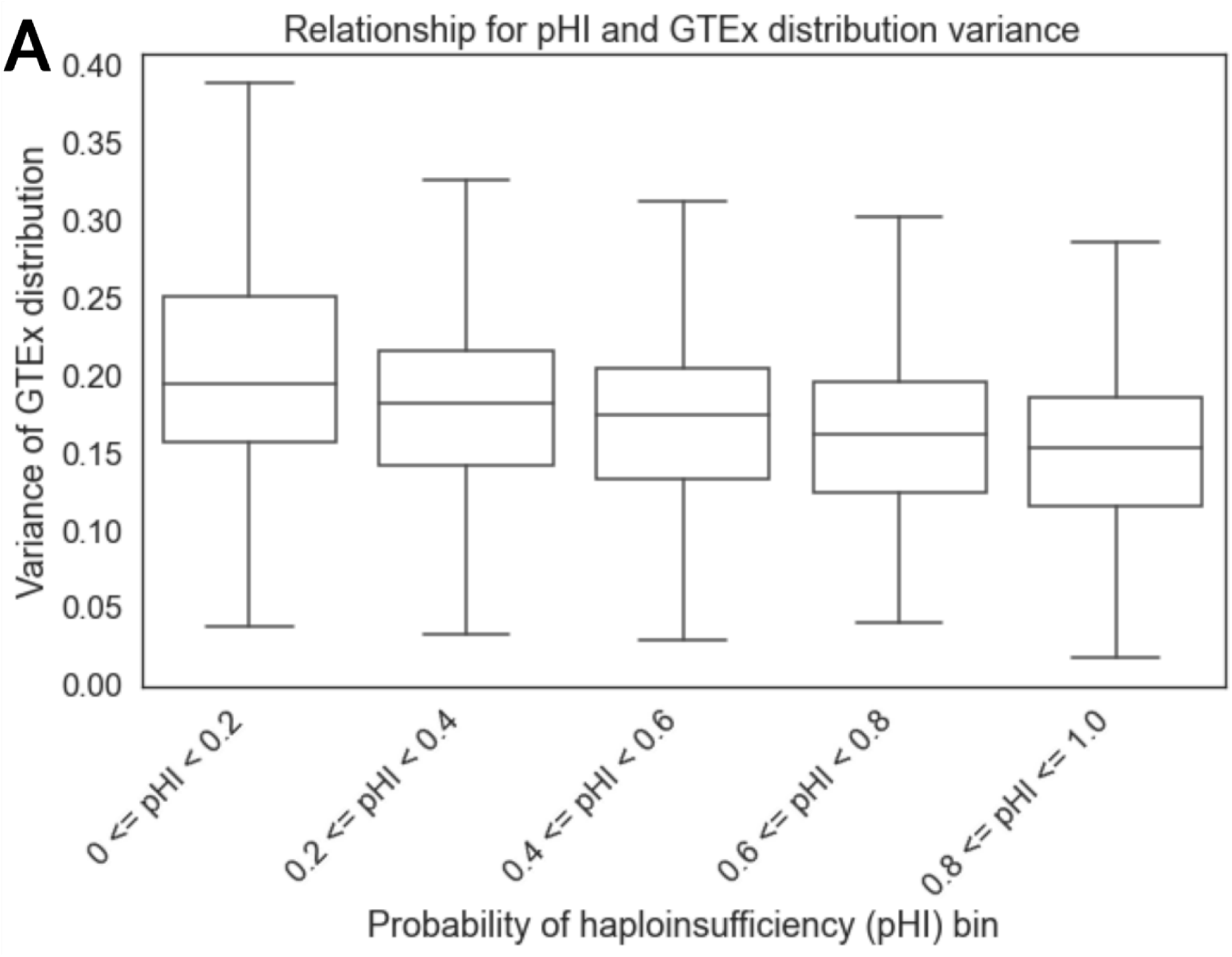
Relationship between the variance of the GTEx ASE distribution and the probability of Haploinsufficiency score: A) Shows the relationship between the variance of the GTEx ASE distribution and the probability of Haploinsufficiency score (a measure of constraint on gene expression) for each gene tested in this manuscript. As the probability of Haploinsufficiency increases, the variance of the GTEx distribution decreases. The spearman correlation is -0.28 with p < 10^-170^.

**Supplementary Figure 2.**
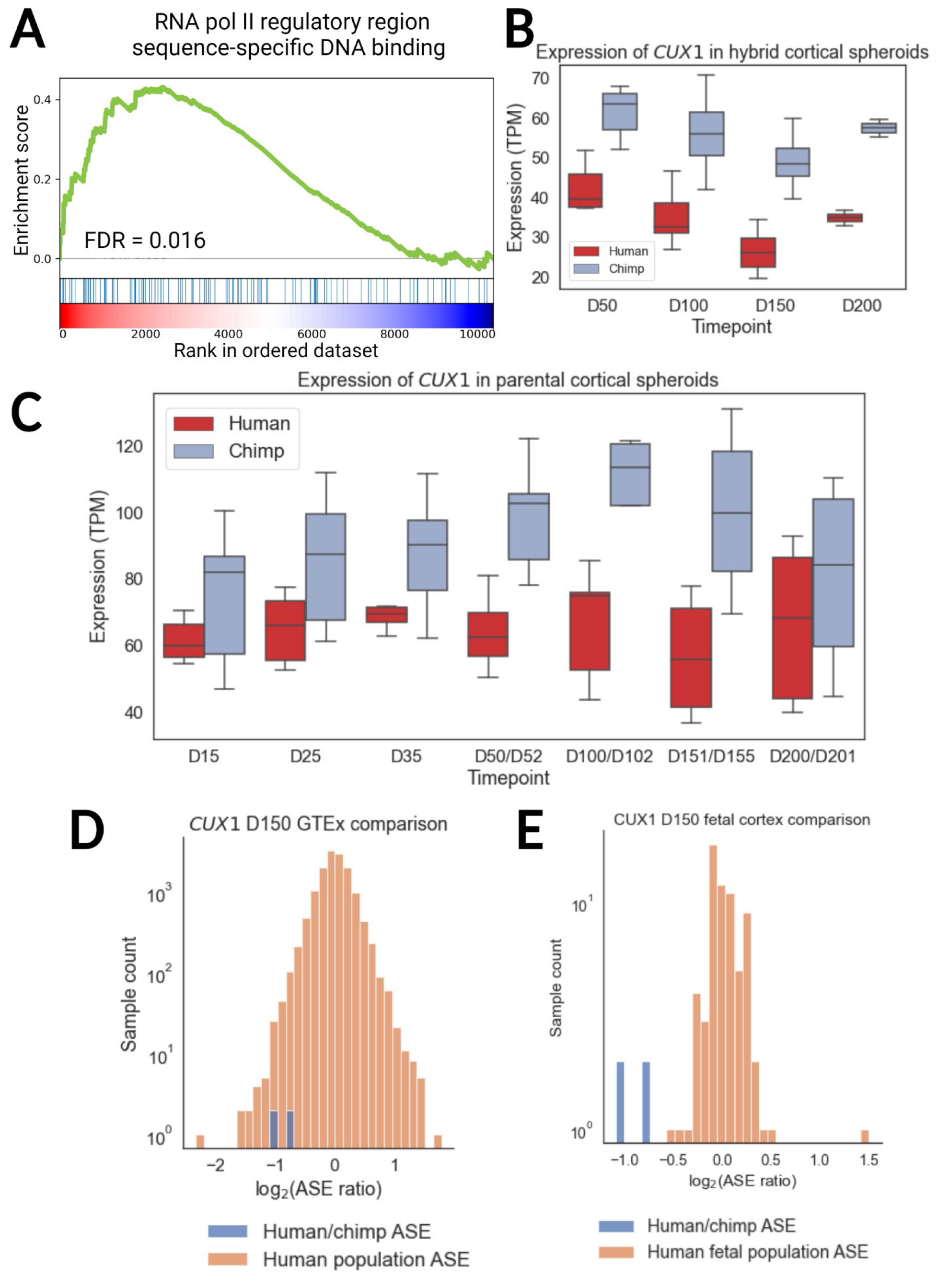
Exploration of changes in CUX1 expression: **A)** Summary of RNA polymerase II regulatory region sequence-specific DNA binding. Each blue line represents a gene in the gene set and the green curve is the cumulative enrichment score. Genes in the gene set are enriched at the top of the list. **B)** Expression of CUX1 in human-chimpanzee hybrid cortical spheroids (CS). Expression is shown in transcripts per million (TPM). Expression from the chimpanzee allele is consistently higher than expression from the human allele. **C)** Expression of CUX1 in human and chimpanzee parental cortical spheroids (CS). Expression is shown in transcripts per million (TPM). **D)** Comparison of the GTEx ASE distribution for CUX1 to the human-chimpanzee hybrid ASE distribution. **E)** Comparison of the fetal cortex ASE distribution for CUX1 to the human-chimpanzee ASE distribution. The human-chimpanzee hybrid ASE values lie well outside the human fetal cortex distribution.

**Supplementary Figure 3.**
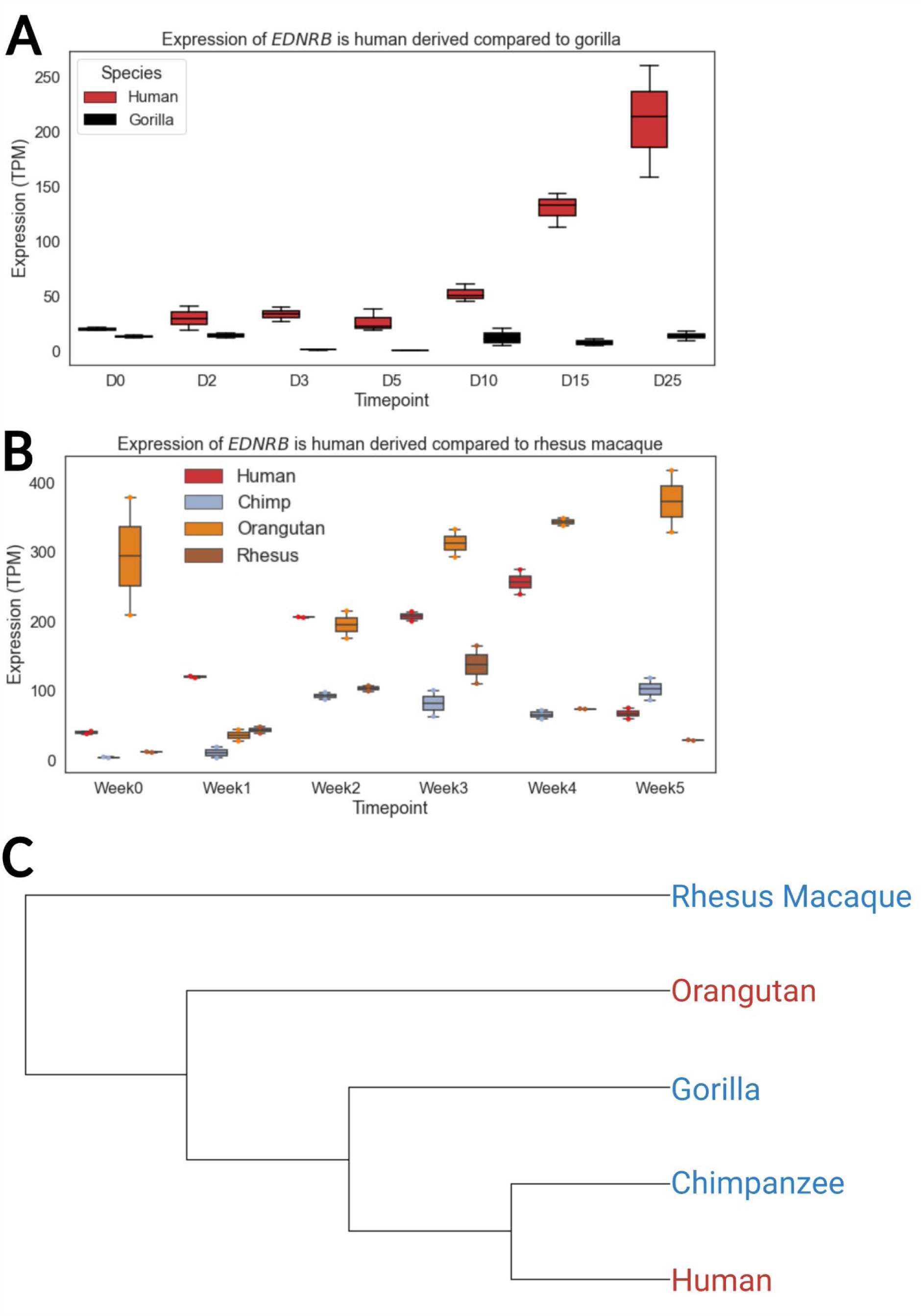
Changes in EDNRB expression are human derived: **A)** Comparison of EDNRB expression between early-stage human and gorilla cerebral organoids [55]. Human expression is considerably higher than gorilla expression across timepoints indicating that EDRNB is upregulated in the human lineage (as opposed to downregulated in the chimpanzee lineage). **B)** Comparison of EDNRB expression between early-stage human, chimpanzee, orangutan, and rhesus macaque cerebral organoids [53]. Human expression is considerably higher than rhesus macaque expression across timepoints. However, orangutan EDNRB expression is high as well, indicating an independent increase in expression in the orangutan lineage. **C)** Phylogenetic tree of old-world primates. Red text indicates a high *EDNRB* expression and blue text indicates low expression. Notably, as gorillas are more closely related to humans than organgutans this implies that the most parsimonious explanation for the data is that the last common ancestor of gorillas, chimpanzees, and humans had low *EDNRB* expression and that there was an increase in the human lineage.

**Supplementary Figure 4.**
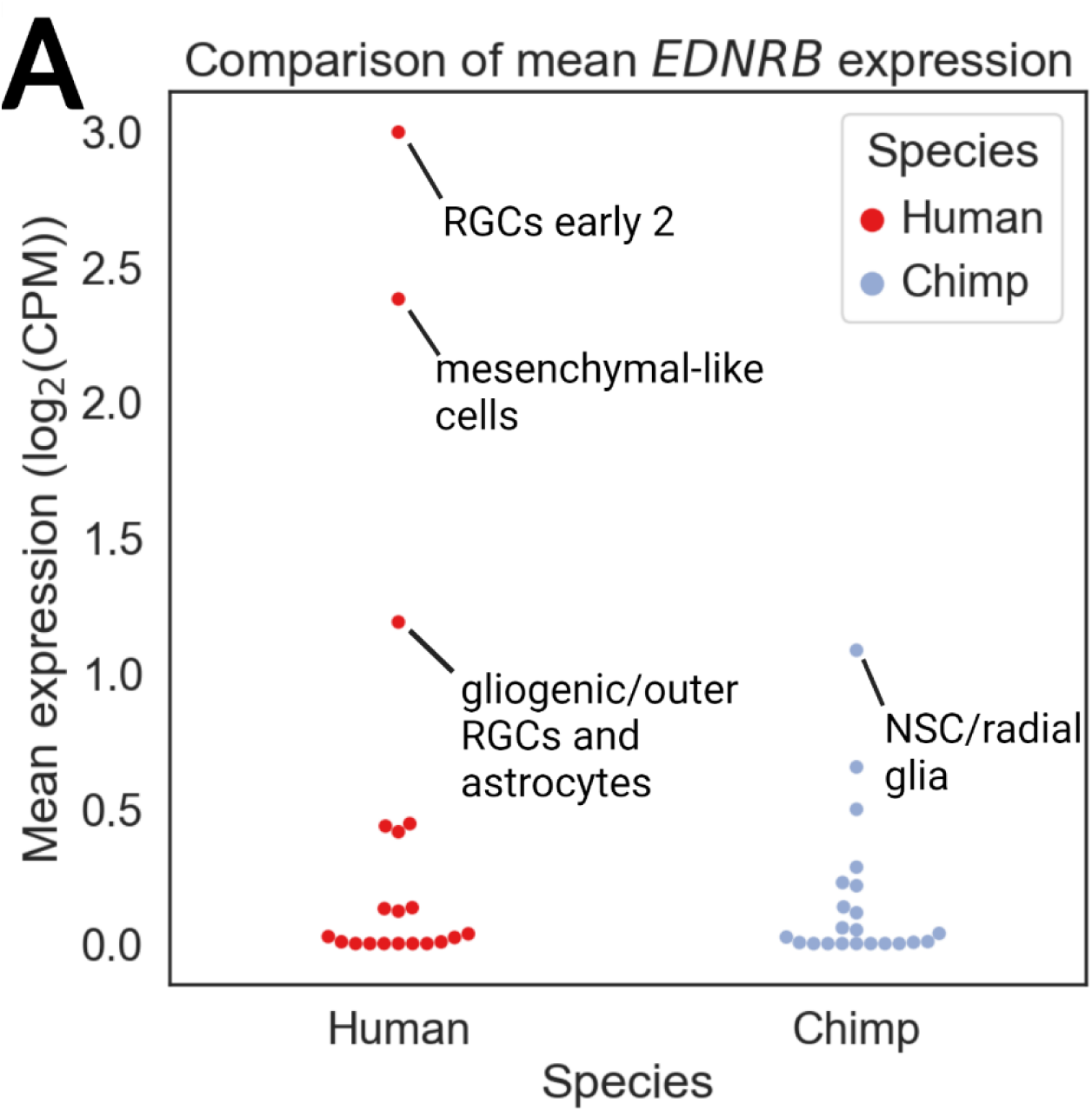
A population of human radial glia expresses EDNRB: **A)** Plot showing mean expression of EDNRB across chimpanzee and human cell clusters [42]. Expression in the “RGCs early 2” cluster is significantly higher than in all chimpanzee clusters by Mann-Whitney U Test (p < 10^-16^ for all comparisons).

**Supplementary Figure 5.**
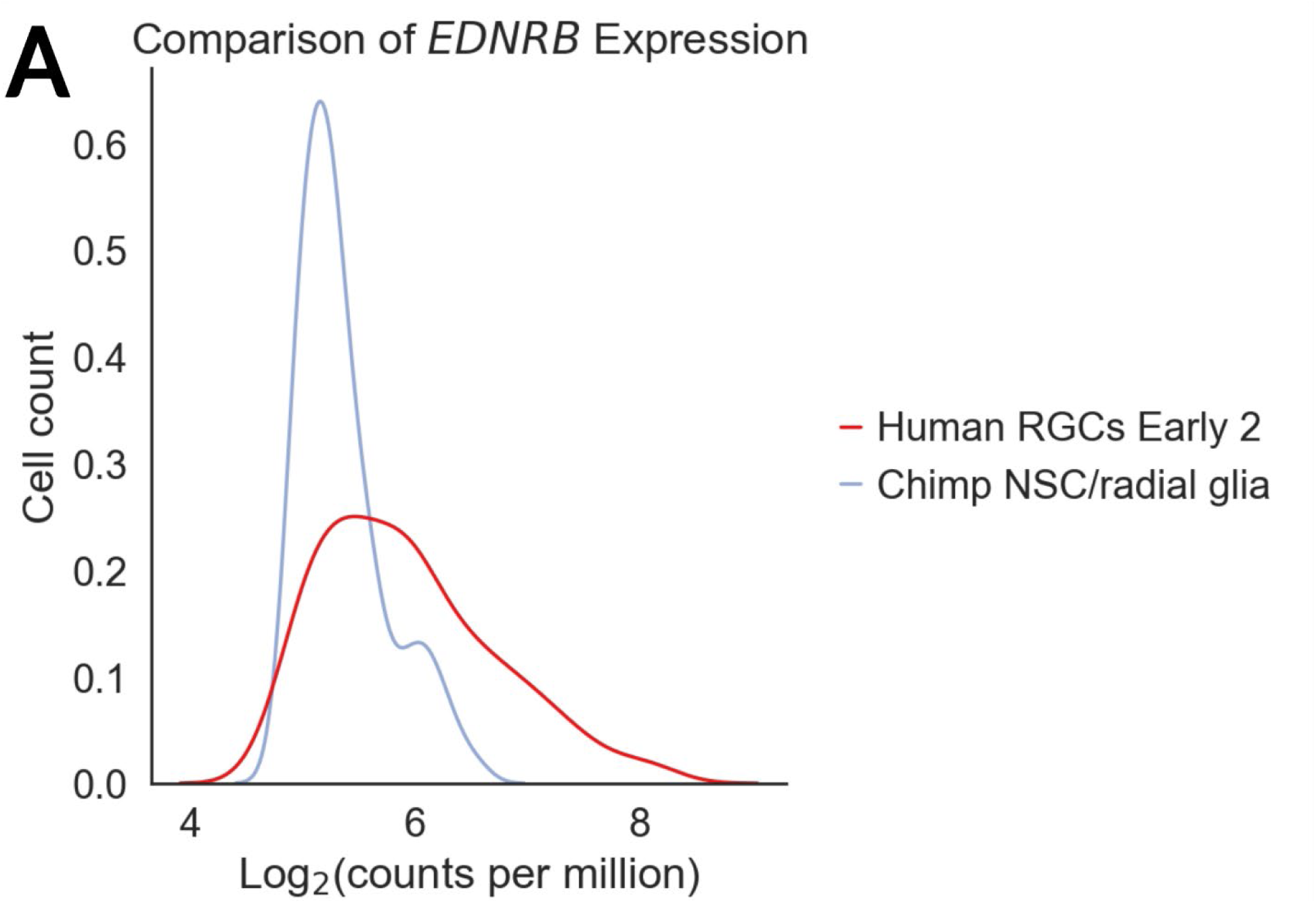
Expression of EDNRB is higher in human radial glia: The distribution of EDNRB expression in cells with non-zero counts [42]. As there were many more chimpanzee cells in the Chimp NSC/radial glia category, we down-sampled the number of cells so that there were an equal number of EDNRB-expressing cells. The distribution in chimpanzee NSC/radial glia is much more shifted left than that of the human “RGCs Early 2” cluster indicating higher expression in the human cluster.

**Supplementary Figure 6.**
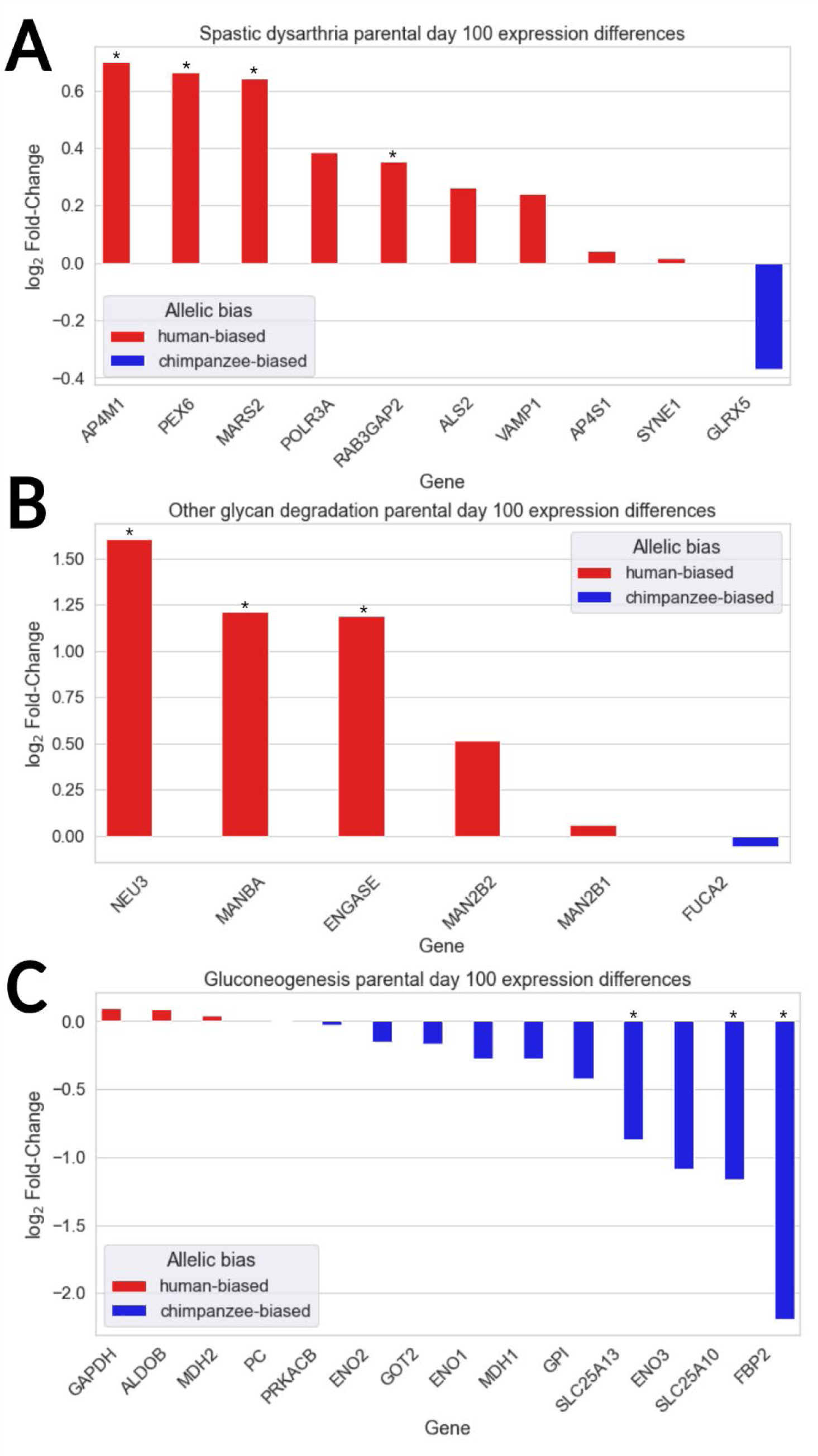
Many gene expression changes have the same direction in hybrid and parental organoids: **A**) Barplot showing the log_2_ fold-changes of genes driving the *Spastic dysarthria* enrichment comparing parental human and chimpanzee cortical spheroids at day 100 of differentiation. Asterisks indicate a significant difference for that gene at and FDR cutoff of 0.1. B) Same as in A but for genes driving the *Other glycan degradation* enrichment. C) Same as in A, but for genes driving the *Gluconeogenesis* enrichment.

**Supplementary Figure 7.**
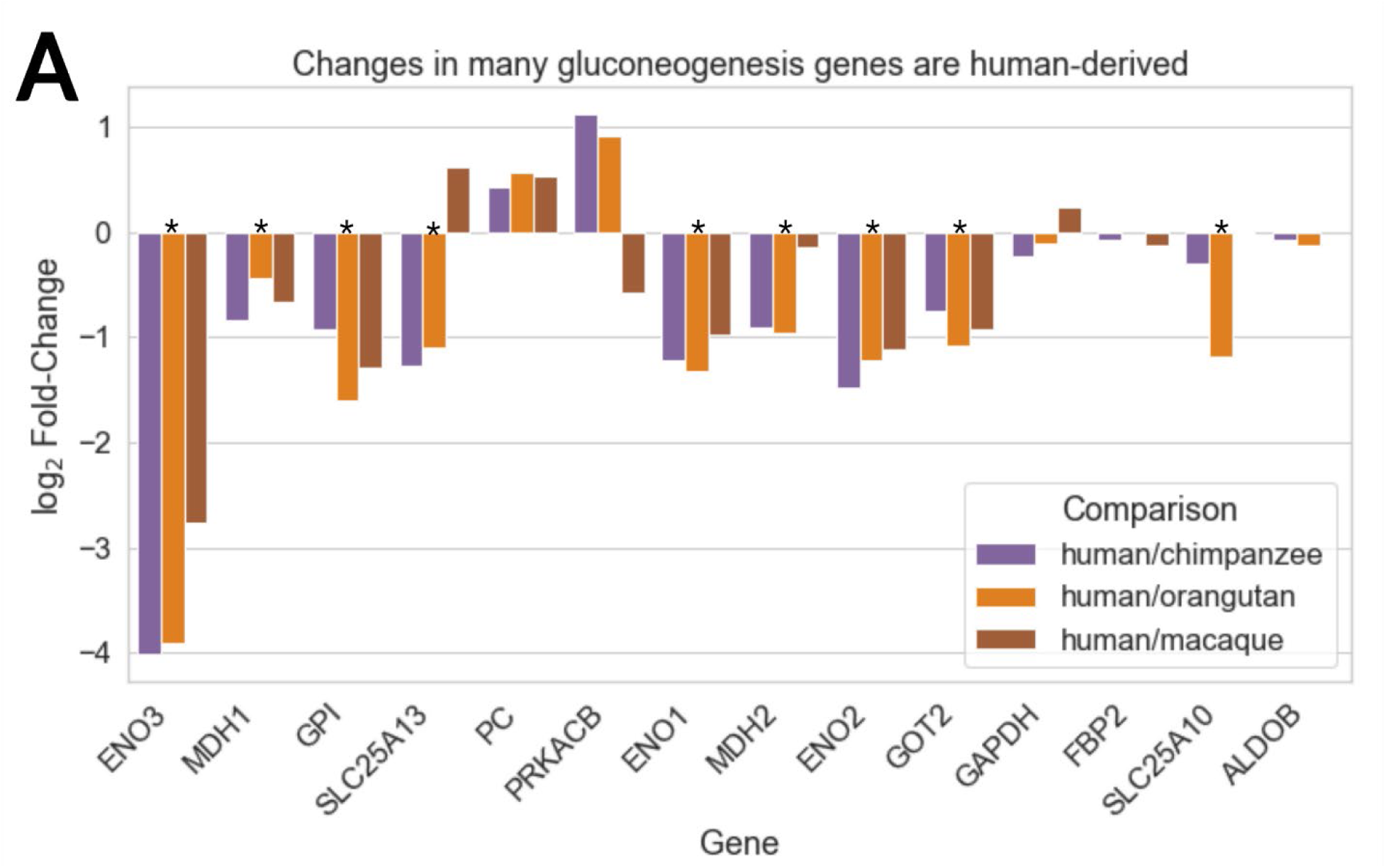
Changes in many gluconeogenesis genes are human-derived: **A**) Barplot showing that many changes in the expression of gluconeogenesis genes are human-derived and occur in parental organoids. Data are from Week 5 cerebral organoids [53]. Asterisks indicate genes for which the human-orangutan difference is significant at an FDR cutoff of 0.1 and with lower expression in human. Genes whose down-regulation is not human-derived generally show insignificant differences in expression between humans and chimpanzees potentially indicating compensatory trans-acting genetic changes.

**Supplementary Figure 9:**
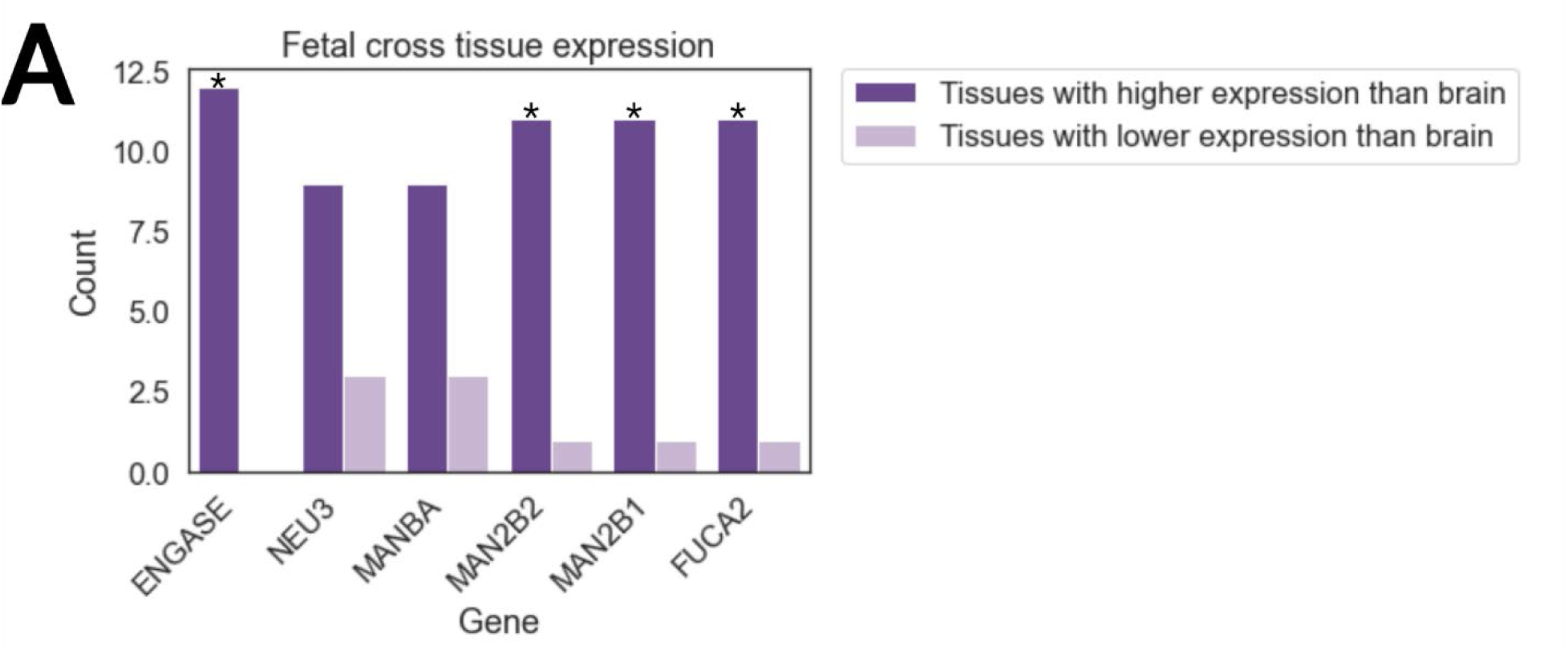
**A**) Explanation of computation of difference in ranks. Population ASE Ranking was subtracted from the traditional ranking. Due to this, genes that are highly ranked in both the traditional and population ASE rankings (likely those with low p-values and high difference in ASE) are near the middle of the list in the difference of ranks. On the other hand, genes with moderately high p-values and high constraint on expression are ranked lowly in the traditional ranking, but near the middle in the population ASE ranking, and so are very high in the difference in ranks.

## References

1. Reilly SK, Noonan JP. Evolution of Gene Regulation in Humans. Annual Review of Genomics and Human Genetics. 2016;17:45–67.

2. Romero IG, Ruvinsky I, Gilad Y. Comparative studies of gene expression and the evolution of gene regulation. Nature Reviews Genetics. 2012;13:505–16.

3. King M-C, Wilson AC. Evolution at Two Levels in Humans and Chimpanzees. Science (1979). 1975;188:107–16.

4. Fraser HB. Gene expression drives local adaptation in humans. Genome Research. 2013;23:1089–96.

5. Kelley JL, Gilad Y. Effective study design for comparative functional genomics. Nature Reviews Genetics. 2020;21:385–6.

6. Housman G, Gilad Y. Prime time for primate functional genomics. Current Opinion in Genetics & Development. 2020;62:1–7.

7. Zhu Y, Sousa AMM, Gao T, Skarica M, Li M, Santpere G, et al. Spatiotemporal transcriptomic divergence across human and macaque brain development. Science (1979). 2018;362.

8. Gokhman D, Agoglia RM, Kinnebrew M, Gordon W, Sun D, Bajpai VK, et al. Human– chimpanzee fused cells reveal cis-regulatory divergence underlying skeletal evolution. Nature Genetics. 2021;53:467–76.

9. Agoglia RM, Sun D, Birey F, Yoon S-J, Miura Y, Sabatini K, et al. Primate cell fusion disentangles gene regulatory divergence in neurodevelopment. Nature. 2021;592:421–7.

10. Hu CK, York RA, Metz HC, Bedford NL, Fraser HB, Hoekstra HE. cis-Regulatory changes in locomotor genes are associated with the evolution of burrowing behavior. Cell Reports. 2022;38:110360.

11. Mack KL, Campbell P, Nachman MW. Gene regulation and speciation in house mice. Genome Research. 2016;26:451–61.

12. Combs PA, Krupp JJ, Khosla NM, Bua D, Petrov DA, Levine JD, et al. Tissue-Specific cis-Regulatory Divergence Implicates eloF in Inhibiting Interspecies Mating in Drosophila. Current Biology. 2018;28:3969–3975.e3.

13. Zhang X, Borevitz JO. Global Analysis of Allele-Specific Expression in *Arabidopsis thaliana*. Genetics. 2009;182:943–54.

14. Song JHT, Grant RL, Behrens VC, Kučka M, Roberts Kingman GA, Soltys V, et al. Genetic studies of human–chimpanzee divergence using stem cell fusions. Proceedings of the National Academy of Sciences. 2021;118.

15. Prud’homme B, Gompel N, Carroll SB. Emerging principles of regulatory evolution. Proceedings of the National Academy of Sciences. 2007;104:8605–12.

16. Wittkopp PJ, Kalay G. Cis-regulatory elements: molecular mechanisms and evolutionary processes underlying divergence. Nature Reviews Genetics. 2012;13:59–69.

17. Blekhman R, Oshlack A, Chabot AE, Smyth GK, Gilad Y. Gene Regulation in Primates Evolves under Tissue-Specific Selection Pressures. PLoS Genetics. 2008;4:e1000271.

18. Gilad Y, Oshlack A, Smyth GK, Speed TP, White KP. Expression profiling in primates reveals a rapid evolution of human transcription factors. Nature. 2006;440:242–5.

19. Roy S, Wapinski I, Pfiffner J, French C, Socha A, Konieczka J, et al. Arboretum: Reconstruction and analysis of the evolutionary history of condition-specific transcriptional modules. Genome Research. 2013;23:1039–50.

20. Rohlfs R V., Nielsen R. Phylogenetic ANOVA: The Expression Variance and Evolution Model for Quantitative Trait Evolution. Systematic Biology. 2015;64:695–708.

21. Castel SE, Aguet F, Mohammadi P, Aguet F, Anand S, Ardlie KG, et al. A vast resource of allelic expression data spanning human tissues. Genome Biology. 2020;21:234.

22. Benjamini Y, Hochberg Y. Controlling the False Discovery Rate: A Practical and Powerful Approach to Multiple Testing. Journal of the Royal Statistical Society: Series B (Methodological). 1995;57:289–300.

23. Love MI, Huber W, Anders S. Moderated estimation of fold change and dispersion for RNA-seq data with DESeq2. Genome Biology. 2014;15:550.

24. Collins RL, Glessner JT, Porcu E, Niestroj L-M, Ulirsch J, Kellaris G, et al. A cross-disorder dosage sensitivity map of the human genome. Genomic Medicine Institute [Internet]. Lerner Research Institute; 15:23–5. Available from: https://doi.org/10.1101/2021.01.26.21250098

25. Paşca AM, Sloan SA, Clarke LE, Tian Y, Makinson CD, Huber N, et al. Functional cortical neurons and astrocytes from human pluripotent stem cells in 3D culture. Nature Methods. 2015;12:671–8.

26. Ferraro NM, Strober BJ, Einson J, Abell NS, Aguet F, Barbeira AN, et al. Transcriptomic signatures across human tissues identify functional rare genetic variation. Science (1979). 2020;369:eaaz5900.

27. Aygün N, Elwell AL, Liang D, Lafferty MJ, Cheek KE, Courtney KP, et al. Brain-trait-associated variants impact cell-type-specific gene regulation during neurogenesis. The American Journal of Human Genetics. 2021;108:1647–68.

28. Subramanian A, Tamayo P, Mootha VK, Mukherjee S, Ebert BL, Gillette MA, et al. Gene set enrichment analysis: A knowledge-based approach for interpreting genome-wide expression profiles. Proceedings of the National Academy of Sciences. 2005;102:15545–50.

29. Platzer K, Cogné B, Hague J, Marcelis CL, Mitter D, Oberndorff K, et al. Haploinsufficiency of *CUX1* Causes Nonsyndromic Global Developmental Delay With Possible Catch-up Development. Annals of Neurology. 2018;84:200–7.

30. Paciorkowski AR, Traylor RN, Rosenfeld JA, Hoover JM, Harris CJ, Winter S, et al. MEF2C Haploinsufficiency features consistent hyperkinesis, variable epilepsy, and has a role in dorsal and ventral neuronal developmental pathways. neurogenetics. 2013;14:99–111.

31. Runge K, Mathieu R, Bugeon S, Lafi S, Beurrier C, Sahu S, et al. Disruption of NEUROD2 causes a neurodevelopmental syndrome with autistic features via cell-autonomous defects in forebrain glutamatergic neurons. Molecular Psychiatry. 2021;26:6125–48.

32. Doan RN, Bae B-I, Cubelos B, Chang C, Hossain AA, Al-Saad S, et al. Mutations in Human Accelerated Regions Disrupt Cognition and Social Behavior. Cell. 2016;167:341–354.e12.

33. Köhler S, Gargano M, Matentzoglu N, Carmody LC, Lewis-Smith D, Vasilevsky NA, et al. The Human Phenotype Ontology in 2021. Nucleic Acids Research. 2021;49:D1207–17.

34. Carbon S, Ireland A, Mungall CJ, Shu S, Marshall B, Lewis S. AmiGO: online access to ontology and annotation data. Bioinformatics. 2009;25:288–9.

35. Jassal B, Matthews L, Viteri G, Gong C, Lorente P, Fabregat A, et al. The reactome pathway knowledgebase. Nucleic Acids Research. 2019;

36. Kanehisa M, Sato Y, Kawashima M, Furumichi M, Tanabe M. KEGG as a reference resource for gene and protein annotation. Nucleic Acids Research. 2016;44:D457–62.

37. Köhler S, Doelken SC, Mungall CJ, Bauer S, Firth H V., Bailleul-Forestier I, et al. The Human Phenotype Ontology project: linking molecular biology and disease through phenotype data. Nucleic Acids Research. 2014;42:D966–74.

38. Girskis KM, Stergachis AB, DeGennaro EM, Doan RN, Qian X, Johnson MB, et al. Rewiring of human neurodevelopmental gene regulatory programs by human accelerated regions. Neuron. 2021;109:3239–3251.e7.

39. Kita R, Venkataram S, Zhou Y, Fraser HB. High-resolution mapping of *cis* -regulatory variation in budding yeast. Proceedings of the National Academy of Sciences. 2017;114:E10736–44.

40. Fraser HB. Genome-wide approaches to the study of adaptive gene expression evolution. BioEssays. 2011;33:469–77.

41. Adams KL, Riparini G, Banerjee P, Breur M, Bugiani M, Gallo V. Endothelin-1 signaling maintains glial progenitor proliferation in the postnatal subventricular zone. Nature Communications. 2020;11:2138.

42. Kanton S, Boyle MJ, He Z, Santel M, Weigert A, Sanchís-Calleja F, et al. Organoid single-cell genomic atlas uncovers human-specific features of brain development. Nature. 2019;574:418–22.

43. Vidovic M, Chen M-M, Lu Q-Y, Kalloniatis KF, Martin BM, Tan AHY, et al. Deficiency in Endothelin Receptor B Reduces Proliferation of Neuronal Progenitors and Increases Apoptosis in Postnatal Rat Cerebellum. Cellular and Molecular Neurobiology. 2008;28:1129–38.

44. Shinohara H, Udagawa J, Morishita R, Ueda H, Otani H, Semba R, et al. Gi2 Signaling Enhances Proliferation of Neural Progenitor Cells in the Developing Brain. Journal of Biological Chemistry. 2004;279:41141–8.

45. Eichmüller OL, Corsini NS, Vértesy Á, Morassut I, Scholl T, Gruber V-E, et al. Amplification of human interneuron progenitors promotes brain tumors and neurological defects. Science (1979). 2022;375.

46. Gkini V, Namba T. Glutaminolysis and the Control of Neural Progenitors in Neocortical Development and Evolution. The Neuroscientist. 2022;107385842110690.

47. Clark HM, Duffy JR, Whitwell JL, Ahlskog JE, Sorenson EJ, Josephs KA. Clinical and imaging characterization of progressive spastic dysarthria. European Journal of Neurology. 2014;21:368–76.

48. Alkhayat AH, Kraemer SA, Leipprandt JR, Macek M, Kleijer WJ, Friderici KH. Human - Mannosidase cDNA Characterization and First Identification of a Mutation Associated with Human -Mannosidosis. Human Molecular Genetics. 1998;7:75–83.

49. Cathey SS, Sarasua SM, Simensen R, Pietris K, Kimbrell G, Sillence D, et al. Intellectual functioning in alpha-mannosidosis. JIMD Reports. 2019;50:44–9.

50. Blomqvist M, Smeland MF, Lindgren J, Sikora P, Riise Stensland HMF, Asin-Cayuela J. β- Mannosidosis caused by a novel homozygous intragenic inverted duplication in *MANBA*. Molecular Case Studies. 2019;5:a003954.

51. Hudson RR, Kreitman M, Aguadé M. A Test of Neutral Molecular Evolution Based on Nucleotide Data. Genetics. 1987;116:153–9.

52. Zhang H, Zhang F, Yu Y, Feng L, Jia J, Liu B, et al. A Comprehensive Online Database for Exploring ∼20,000 Public Arabidopsis RNA-Seq Libraries. Molecular Plant. 2020;13:1231–3.

53. Field AR, Jacobs FMJ, Fiddes IT, Phillips APR, Reyes-Ortiz AM, LaMontagne E, et al. Structurally Conserved Primate LncRNAs Are Transiently Expressed during Human Cortical Differentiation and Influence Cell-Type-Specific Genes. Stem Cell Reports. 2019;12:245–57.

54. Zhu Y, Sousa AMM, Gao T, Skarica M, Li M, Santpere G, et al. Spatiotemporal transcriptomic divergence across human and macaque brain development. Science (1979). 2018;362.

55. Benito-Kwiecinski S, Giandomenico SL, Sutcliffe M, Riis ES, Freire-Pritchett P, Kelava I, et al. An early cell shape transition drives evolutionary expansion of the human forebrain. Cell. 2021;184:2084–2102.e19.

56. Khrameeva E, Kurochkin I, Han D, Guijarro P, Kanton S, Santel M, et al. Single-cell-resolution transcriptome map of human, chimpanzee, bonobo, and macaque brains. Genome Research. 2020;30:776–89.

57. Dobin A, Davis CA, Schlesinger F, Drenkow J, Zaleski C, Jha S, et al. STAR: ultrafast universal RNA-seq aligner. Bioinformatics. 2013;29:15–21.

58. Broad Institute. Picard Toolkit. 2019.

59. Givanna H Putri SAPTPJEPFZ. Analysing high-throughput sequencing data in Python with HTSeq 2.0. ArXiv. 2021;

60. Li B, Ruotti V, Stewart RM, Thomson JA, Dewey CN. RNA-Seq gene expression estimation with read mapping uncertainty. Bioinformatics. 2010;26:493–500.

61. Zhu A, Ibrahim JG, Love MI. Heavy-tailed prior distributions for sequence count data: removing the noise and preserving large differences. Bioinformatics. 2019;35:2084–92.

62. Supek F, Bošnjak M, Škunca N, Šmuc T. REVIGO Summarizes and Visualizes Long Lists of Gene Ontology Terms. PLoS ONE. 2011;6:e21800.

63. Wolf FA, Angerer P, Theis FJ. SCANPY: large-scale single-cell gene expression data analysis. Genome Biology. 2018;19:15.

64. Cao J, O’Day DR, Pliner HA, Kingsley PD, Deng M, Daza RM, et al. A human cell atlas of fetal gene expression. Science (1979). 2020;370.

